# Network reconstruction and modelling made reproducible with moped

**DOI:** 10.1101/2020.12.04.411512

**Authors:** Nima P Saadat, Marvin van Aalst, Oliver Ebenhöh

## Abstract

Mathematical modeling of metabolic networks is a powerful approach to investigate the underlying principles of metabolism and growth. Such approaches include, amongst others, differential equation based modeling of metabolic systems, constraint based modeling and topological analysis of metabolic networks. Most of these methods are well established and are implemented in numerous software packages, but these are scattered between different programming languages, packages and syntaxes. This complicates establishing straight forward pipelines integrating model construction and simulation. We present the Python package moped which serves as an integrative hub for reproducible construction, modification, curation and analysis of metabolic models. moped supports draft reconstruction of models directly from genome/proteome sequences and pathway/genome databases utilizing GPR annotations, providing a completely reproducible model construction and curation process within executable Python scripts. Alternatively, existing models published in SBML format can be easily imported. Models are represented as Python objects, for which a wide spectrum of easy-to-use modification and analysis methods exist. The model structure can be manually altered by adding, removing or modifying reactions, and gap filling reactions can be found and inspected. This greatly supports the development of draft models, as well as the curation and testing of models. Moreover, moped provides several analysis methods, in particular including the calculation of biosynthetic capacities using metabolic network expansion. The integration with other Python based tools is facilitated through various model export options. For example, a model can be directly converted into a CobraPy object for constraint-based analyses. moped is a fully documented and expandable Python package. We demonstrate the capability to serve as a hub for integrating reproducible model construction and curation, database import, topological analysis and export for constraint-based analyses.

## Introduction

Theoretical analysis of metabolic pathways has a long standing tradition. The early approaches to study glycolosis, for example, have considerably increased our understanding of fundamental regulatory principles in metabolism [1]. In recent approaches, metabolic modelling was employed to study metabolic interdependencies in microbial communities, and to identify putative drug targets for microbial pathogens [2,3].

Several theoretical techniques to study metabolism have been established. The most classic technique is the analysis of metabolic networks by representing them as systems of ordinary differential equations (ODEs). This representation heavily depends on detailed knowledge of stoichiometries, parameters of enzyme kinetics and regulatory mechanisms of reactions [4]. This approach is extremely useful for investigating relatively small systems. The upsurge of novel high-throughput experimental ‘omics’ techniques led to the collection of immense amounts of data, resulting in an ever increasing number of fully sequenced genomes. The improved quality of annotated genes resulted in a tremendous increase in information of enzymes and the respective metabolic reactions. This information has been collected in biochemical databases like MetaCyc, BioCyc, KEGG or BiGG [5–8]. Such databases provide information for large-scale metabolic networks of many different organisms. However, analysing such large-scale metabolic networks using systems of ordinary differential equations is difficult. This is, to a large extent, due to missing information on kinetic parameters of the involved enzymatic reactions [9]. One convenient alternative is constraint-based modelling, and its mathematical method flux balance analysis (FBA) [10]. This commonly used approach uses the stoichiometric matrix of a reaction network and finds a steady-state vector of reaction fluxes that maximizes or minimizes an objective function that linearly depends on the reaction rates. Other structural analysis techniques focus on the topology of metabolic networks [11]. One such technique is metabolic network expansion and the related concept of metabolic scopes. The metabolic scope describes the set of metabolites, which are topologically producible by a given network from an initial set of compounds [12]. Thus, metabolic network expansion allows to functionally characterize metabolic networks with respect to their biosynthetic capacities [13].

Topological techniques are extremely useful in the process of curating models, in particular to identify and add missing reactions [14]. This process, called gap filling, allows, for example, to complement draft metabolic networks in order to guarantee that observed compounds can be produced from the growth medium [15].

Many of the techniques described above have been implemented as Python packages. However, most of these software packages are not directly compatible with each other.

In this work, we present `moped`, a compact but very useful Python package that serves as a hub, offering tools for analysis, development and extension or modification of metabolic models. The integration of BLAST and pathway/genome databases such as MetaCyc and BioCyc into `moped` furthermore allows reconstructing metabolic network models directly from genome sequences [16], and ensuring that the reconstruction process is fully transparent and reproducible. In addition to the de novo construction of models, `moped` provides an interface to import existing metabolic network models in SBML format.

To facilitate curation of metabolic models, `moped` provides an interface to Meneco, a topological gap-filling tool based on answer set programming [17]. All models created with `moped` can easily be exported as CobraPy objects, thus directly integrating constraint-based with model construction and modification [18]. It is even possible to extract a scaffold model of metabolic pathways for kinetic modelling via `modelbase` [19]. The Python package `moped` presented here is the first mathematical modelling hub, which allows constructing reproducible metabolic models de novo, integrating existing models in SMBL format, curating models by gap-filling, and performing topological or constraint-based analyses.

## Implementation

### General Implementation and structure of Python object/classes

`moped` is a package fully integrated in the object-oriented programming language Python. The core is the `moped.Model` class, which instantiates a metabolic network from scratch or from input files like SBML or pathway/genome database (PGDB) flat files. This class includes reactions and compounds, which contain all extracted information for the respective attributes of the metabolic network, which is mostly stored in dictionary structures. The `moped.databases.Cyc` class builds the `moped.Model` object by parsing PGDBs.

The `moped.analysis` module includes the module `moped.analysis.blast`, which constructs a `moped.Model` from all reactions in the MetaCyc database that can be found within genome/proteome sequences using BLAST and GPR rules. For this, `moped` requires FASTA files as input, and parameters for BLAST can be specified.

The module `moped.analysis.gapfilling` allows gap-filling via Meneco using one `moped.Model` object as the draft network and another one as the repair network. The last module, `model.analysis.scope` provides topological analysis of `moped.Model` objects using metabolic network expansion. The core classes are displayed in an unified modelling language graph in Figure 2. `moped` can be installed using `pip install moped==1.6.5`.

**Figure 1.**
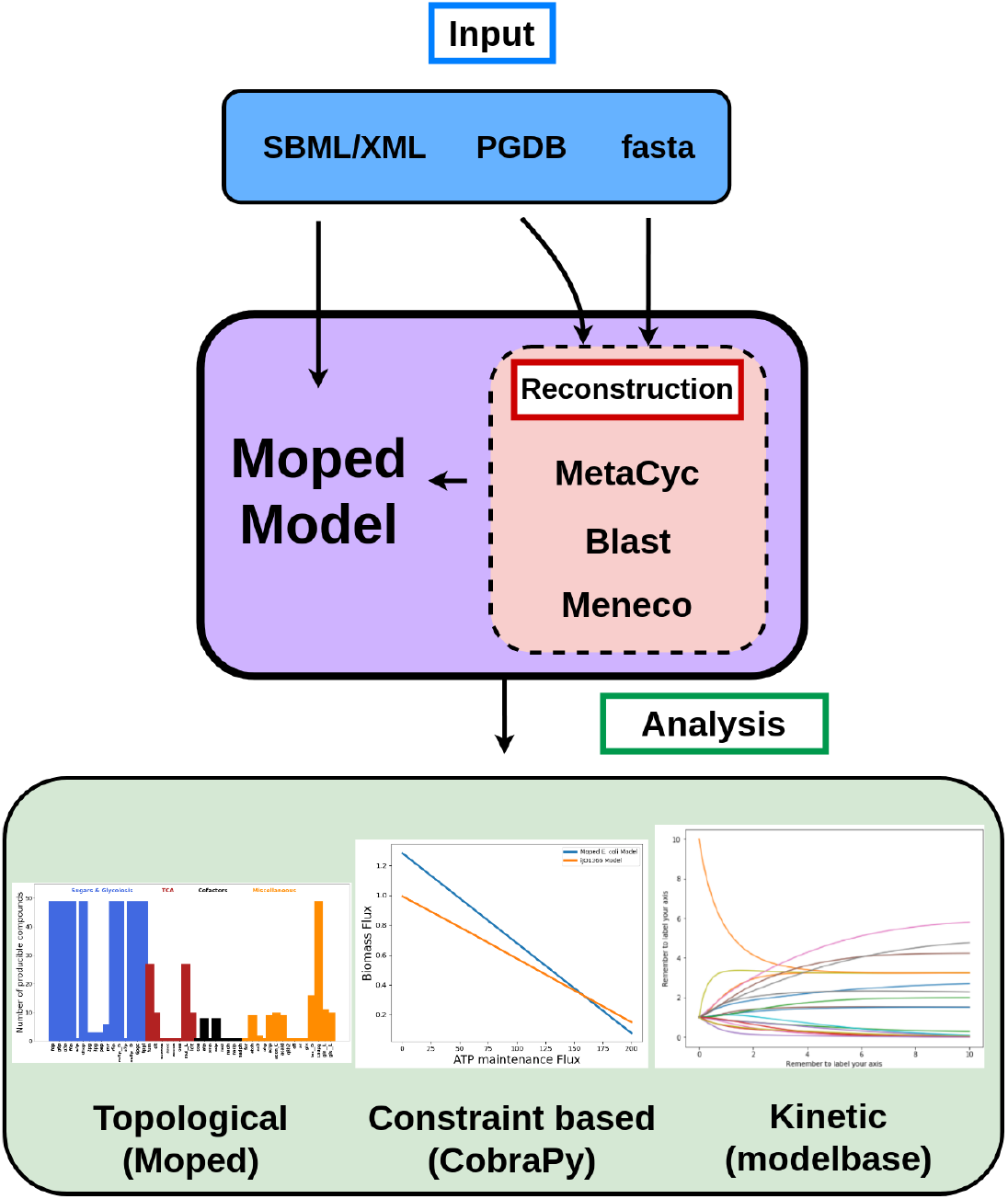
The modelling hub `moped`. `moped` accepts SBML, FASTA files or MetaCyc and BioCyc PGDBs as inputs. PGDBs and SBML files are directly converted into a `moped` object. By BLASTing genome/proteome-sequences against MetaCyc, `moped` models can be constructed utilizing GPR rules. Further reconstruction can be achieved using Meneco for gap-filling. Topological model analysis is implemented in `moped`. For contraint based and kinetic analysis, `moped` offers export as CobraPy and `modelbase` objects, respectively [18,19]

**Figure 2.**
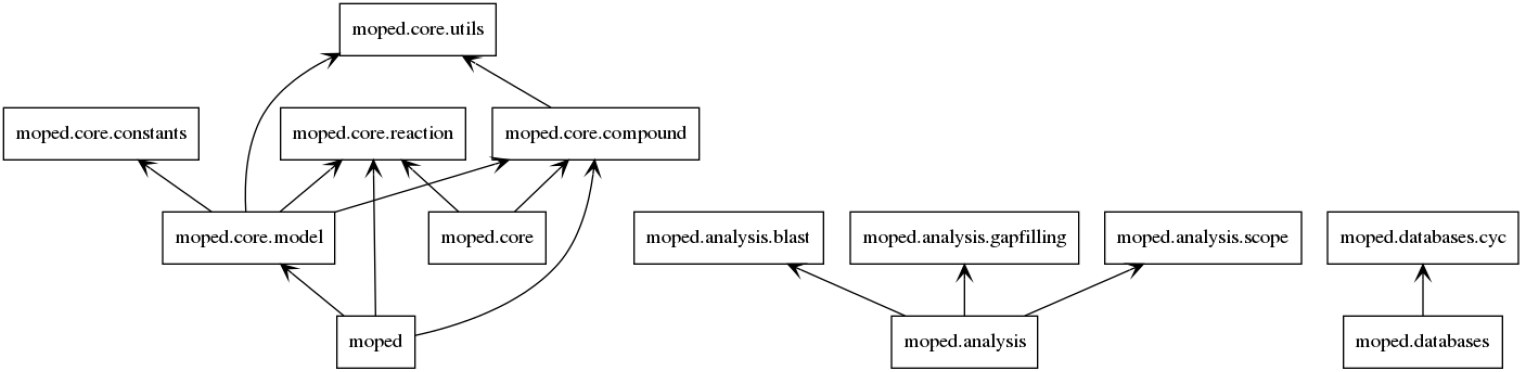
Core classes in `moped`. Details of object attributes and methods can be found in the documentation.

### Model import, extension and modification

`moped` uses SBML files or PGDB flat files as input for constructing a metabolic network model. PGDBs are organism specific pathway/genome databases containing predicted reactions and compounds of the metabolism of the organism [6]. These databases further include detailed information about reactions and compounds, such as sum formulas, charges, references to other database entries or subcellular localisation. This information is of great importance for a consistent analysis of metabolic networks. SBML files represent metabolic networks in an XML-based format and can be considered as a standard for the exchange of reconstructed and curated metabolic models between tools and platforms [20]. Such files can be, amongst others, obtained from databases like BiGG, which provides SBML files of curated metabolic models together with information about the corresponding publications [7].

Because of the wide range of import methods (FASTA, PGDBs and SBML), one particular strength of `moped` is the integration of several analysis tools. Furthermore, `moped` provides a very accessible environment to extend or modify constructed or imported models. Therefore, adding alternative or additional metabolic pathways to pre-existing models, as well as modifying compound and reaction identifiers, is simple and straight forward. Naturally, all `moped` objects can be exported as SBML.

### Metabolic network expansion and tools for topological analysis

A useful and valuable feature of `moped` is the fully implemented network expansion algorithm to perform topological analyses on `moped` objects. Metabolic network expansion can be used to investigate structural properties of metabolic networks, such as biosynthetic capacities and their robustness against structural perturbations [12]. The core concept of metabolic network expansion is the metabolic scope, which contains all compounds that are producible by a network from a given initial set of compounds, termed the seed (see Figure 3). In the expansion process, the seed is used to find all reactions that can proceed in their annotated direction. The respective products are then added to the seed, forming the new seed for the next expansion step. This process continues until no new compounds can be added to the seed. Thus, scopes characterise biosynthetic capacities of metabolic networks, based exclusively on their topology.

**Figure 3.**
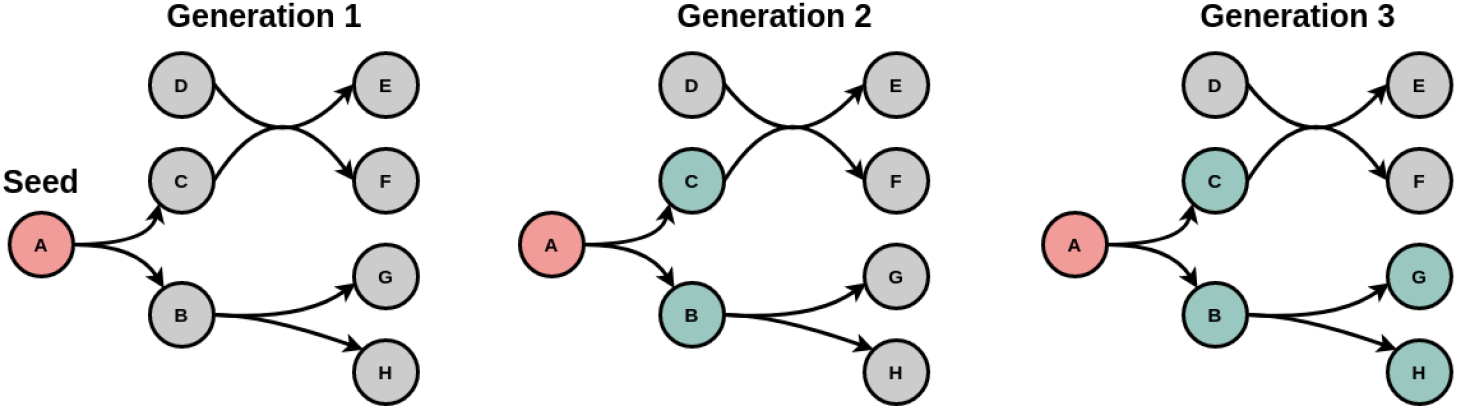
Metabolic Network Expansion Beginning with an initial set of compounds, the seed (here **A**), the expansion process detects all producible compounds in a network and adds them to the seed for the next generation until no additional producible compounds are found.

Any topological analysis based on network expansion depends on a precise definition of reaction reversibilities and involved cofactors. Network expansion uses the stoichiometry of reactions to identify producible compounds. However, stoichiometric coefficients of reactions are annotated for one particular direction. To include the opposite direction (for reversible reactions) into the topological analysis, `moped` provides a method for reversibility duplication. As illustrated in Figure 4 for Triose-phosphoisomerase as an example, this method finds all reversible reactions in a `moped` object and adds the reversed reaction to the network. The new reaction identifier is identical to the identifier of the original reaction concatenated with the suffix ’_rev_’. This model modification can be reverted, if no longer needed.

**Figure 4.**
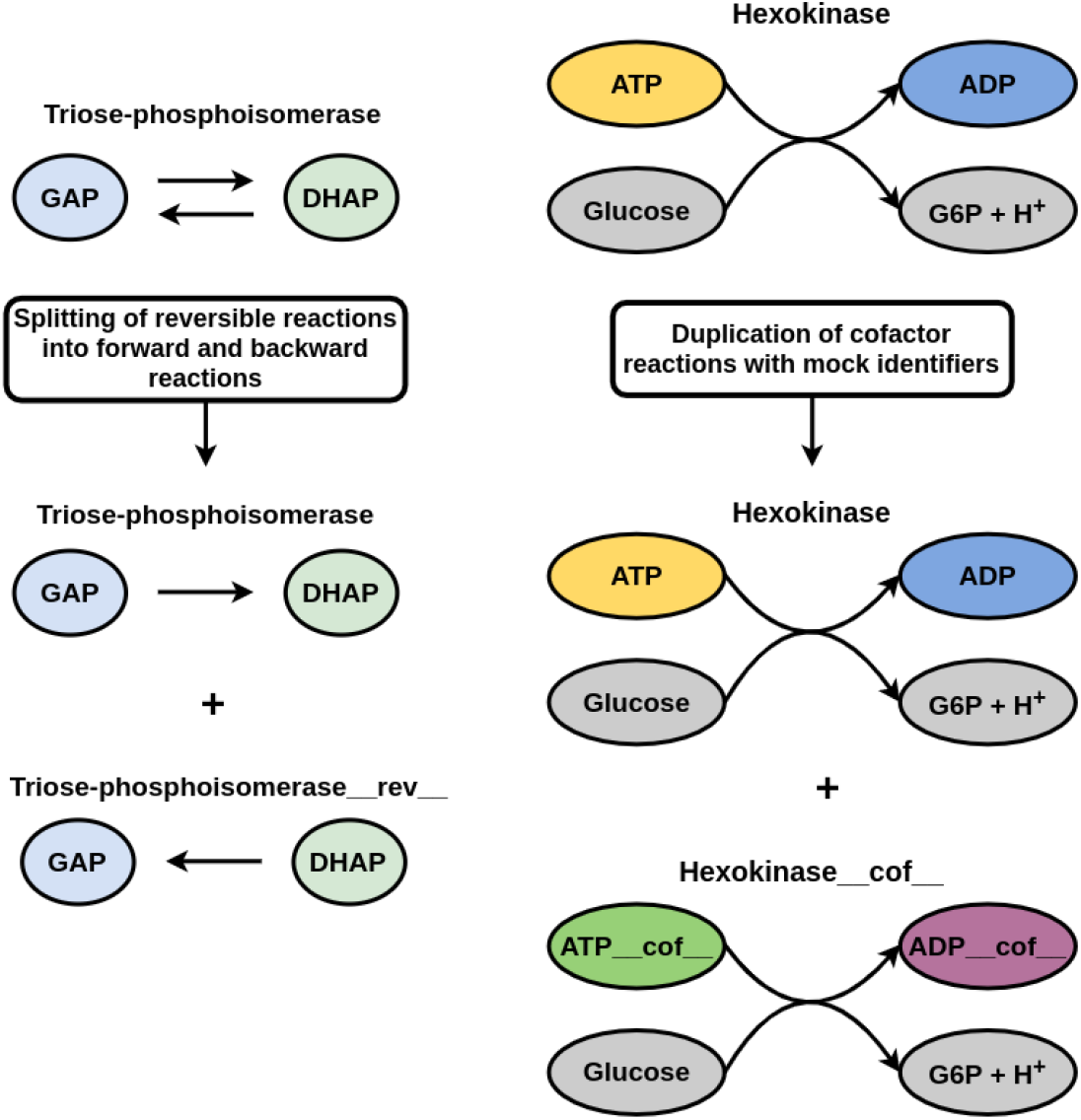
Topological network modifications `moped` offers functions for splitting reversible reactions into forward and backward reactions in a network. Adding a copy of each cofactor dependent reaction and replacing cofactors (here ATP and ADP) with mock identifiers allows unblocking cofactor dependent reactions while avoiding degradation products of cofactors contained in the seed. Such modifications enable biologically feasible topological analysis.

Many reactions depend on specific cofactors. Cofactors usually appear in pairs. One of the most prominent examples is the cofactor pair ATP and ADP. In the majority of reactions with ATP as substrate, ATP serves as a donor of a phosphate group, thus producing ADP. Only a few reactions actually modify the adenosine moiety (for example in nucleotide de novo synthesis). In network expansion, therefore, no reaction utilising ATP or ADP as cofactor could proceed, unless these compounds are either included in the seed or can be produced from metabolites within the seed. If the purpose of network expansion is to realistically calculate a set of producible compounds, this behaviour is not desired, because it leads to a drastic underestimation of the scope. The most naive approach to directly include cofactors in the seed yields misleading results, because in such a case all compounds that can be generated from digesting, e.g., ATP would be included in the scope.

A pragmatic approach to solve this problem is the duplication of cofactors as proposed in [12]. Here, reactions with cofactor pairs are duplicated, where the copied reactions contain “mock cofactors”. In contrast to the real cofactors, the mock cofactors only occur in reactions, in which they act in their role as cofactors. For ATP, this is the transfer of a phosphate group, for NADH or NADPH the transfer of protons and electrons, and for acetyl coenzyme-A, the transfer of the acetyl group. The cofactor duplication allows the use of mock cofactors inside the initial seed. Reactions depending on cofactors might now be able to occur in the expansion process. However, reactions using the cofactors as proper substrates can only occur if the real cofactor can be produced from the seed.

`moped` provides a convenient method for finding and duplicating all reactions using cofactor pairs. The cofactor pairs can either be automatically determined by `moped` for networks imported from BiGG or MetaCyc, or they can be declared individually by the user. The identifiers of the duplicated cofactors are replaced by mock identifiers, which contain the suffix ’_cof_’. The same modification is applied to the respective reaction identifiers. This model modification can be reverted if no longer needed.

The implemented methods for cofactor and reversibility duplication are commonly used to obtain biologically meaningful results for metabolic network expansion. However, they are also highly useful for topological gap-filling using Meneco, during model reconstruction. This is further explained in the next section.

### Reconstruction of draft network models

Construction of metabolic networks highly depends on reliable databases. In order to enable user-friendly metabolic network reconstruction, `moped` includes methods for importing data from the MetaCyc and BioCyc databases, identifying homologous sets of genes with BLAST and gap-filling.

MetaCyc is a universal, highly curated reference database of metabolic pathways and biochemical reactions from all domains of life. BioCyc is a database of organism specific PGDBs containing metabolic network information based on predictions by the PathoLogic component of the Pathway Tools software [21,22]. The MetaCyc and BioCyc databases provide many advantages. Both databases are freely available for academic and nonprofit users. All PGDBs are available in useful flat file formats. Furthermore, these databases include information on the reaction directions based on experimental references and thermodynamics, extensive annotations and therefore information about Gene-Protein-Reaction (GPR) associations, as well as thermodynamic information about metabolites and reactions such as the Gibbs energy of formation and the standard Gibbs energy of reactions.

In order to use BioCyc and MetaCyc for metabolic network construction and analysis, `moped` offers a parser for PGDBs, allowing direct construction of `moped` objects from MetaCyc or BioCyc flat files. `moped` objects can directly be used for network analyses including network expansion and constraint-based modelling. Especially for the latter, it is extremely important that all reactions are mass- and charge-balanced to ensure that all solutions obey mass conversation. Therefore, only reactions which are mass- and charge-balanced are parsed in `moped`. While this process has the danger of omitting annotated genes, including reactions that are not mass- or charge-balanced would violate fundamental physical principles and lead to unrealistic model properties. This pipeline provided by the database import and parsing of `moped` makes it straight forward to construct prokaryotic network models. For eukaryotic metabolic networks, however, intensive and careful curation is required due to missing compartment information. More detailed information about the parsing of PGDBs using `moped` can be found in the documentation.

There exist several pipelines to automatically extract a set of metabolic reactions from a genome or proteome sequence. One popular pipeline is the above mentioned PathoLogic software. `moped` integrates such a pipeline into the Python programming language, directly converting a genome/proteome sequence into a `moped` object which can be immediately used for modelling applications. This functionality is provided by an implemented wrapper for local BLAST to find enzyme reactions in genome sequence fasta files or proteome fasta files by similarity search against enzyme reference sequences from the MetaCyc database. This method constructs a new `moped` object representing a metabolic network of all reactions that are found in a genome sequence or proteome using enzyme monomer amino acid sequences and protein-reaction annotations from MetaCyc to ensure fulfilled gene-protein-reaction associations (GPRs) in all found reactions [23]. This process can be controlled by user defined thresholds. This integrated pipeline makes the model reconstruction perfectly reproducible, and illustrates the functionality of `moped` as a modelling hub.

The next curation step after the initial automatic network construction is usually gap-filling. This describes a process in which reactions are added to the network in order to ensure that the reconstructed model reflects experimentally observed behaviour, such as the production of experimentally measured compounds from the growth medium [24]. There are many available gap-filling methods like GapFill or MIRAGE [25,26]. Most of these methods are based on constraint-based approaches. A common problem is that these approaches can predict gap-filling solutions which use thermodynamically infeasible cycles. In this sense, these approaches are sensitive to self-producing or energy generating cycles [17]. Meneco, in contrast, is a topological gap-filling tool based on the network expansion algorithm. Meneco calculates a minimal set of reactions that need to be added to a draft network such that a specified list of target compounds can be produced from a given set of seed compounds. This gap-filling approach offers the advantage that it is inherently impossible for gap-filling solutions to depend on infeasible cycles. Meneco gap-filling can be directly applied as a method to `moped` objects. One `moped` object represents the draft network and a second the repair network, from which the added reactions are chosen.

The topological network modifications, i.e. reversibility and cofactor duplication, harmonize ideally with the application of Meneco, resulting in networks with biologically realistic behaviour. This again illustrates the integrative nature of the modelling hub `moped`. For an accurate manual curation, automatically determined gap-filling reactions should always be manually inspected before adding them to the network model.

A major advantage and distinguished feature of `moped` is the complete reproducibil-ity of the construction of draft models and the subsequent manual curation. In `moped`, the user can add and remove reactions, or even entire pathways, from draft networks. Furthermore, the user can inspect the reactions found by Meneco to fill gaps and decide if these reactions are valid for specific models. All user decisions become part of an executable Python script, making them perfectly reproducible by others. This underlines that in moped early curation can be integrated closely into the draft model reconstruction process. Likewise, for constraint-based modelling, the user can define which exchange reactions are to be included and, if desired, define their own specific objective functions. Reconstructing draft networks in `moped` lays the ground for model curation without the need to change software environments. In all reconstruction and curation steps, user decisions are documented as commands in an executable Python script, thus making them fully reproducible and transparent.

## Results

### Displaying the advantage of cofactor duplications in topological network analysis

To display the benefits of including the `moped` cofactor duplication, three established models of *E. coli, B. subtilis* and *Synechocystis sp*. PCC 6803 have been parsed into `moped` for a comparative topological analysis [27–29]. In this analysis, we calculated all single metabolite scopes (i.e. the scopes for the seed consisting only of a single metabolite and water) for the respective models. This has been done in three variations: i) including no cofactors to the seed, ii) including the original cofactor compounds, and iii) incuding on the mock cofactors resulting from cofactor duplication (see above). Figure 5 displays the scope sizes (number of compounds contained in the scope) for each model and each variant to calculate the scopes. Apparently, without cofactors, the scopes are very small for most compounds (blue lines). This can be explained by the missing connectivity for reactions that require cofactors. The analysis including the actual cofactor compounds in the seed (orange lines) displays an unrealistically large metabolic scope for every compound, even for inorganic metabolites. This can be explained by the fact that cofactors are usually rather complex metabolites, and now all degradation processes are included during the network expansion. Therefore, the resulting metabolic scopes are no longer reflecting the property of the compound of interest, but rather the degradation products of the metabolized cofactor compounds. The corresponding analysis of models using cofactor duplication and mock cofactors duplicates in the seed (green lines) demonstrates that for small or inorganic metabolites, the scope is still relatively small. For more complex organic compounds, the metabolic scope is increasing without artificially increasing the scope size with degradation products of cofactors. This demonstrates the perks of including cofactor duplication and mock cofactors in seeds for biologically more realistic topological analyses.

**Figure 5.**
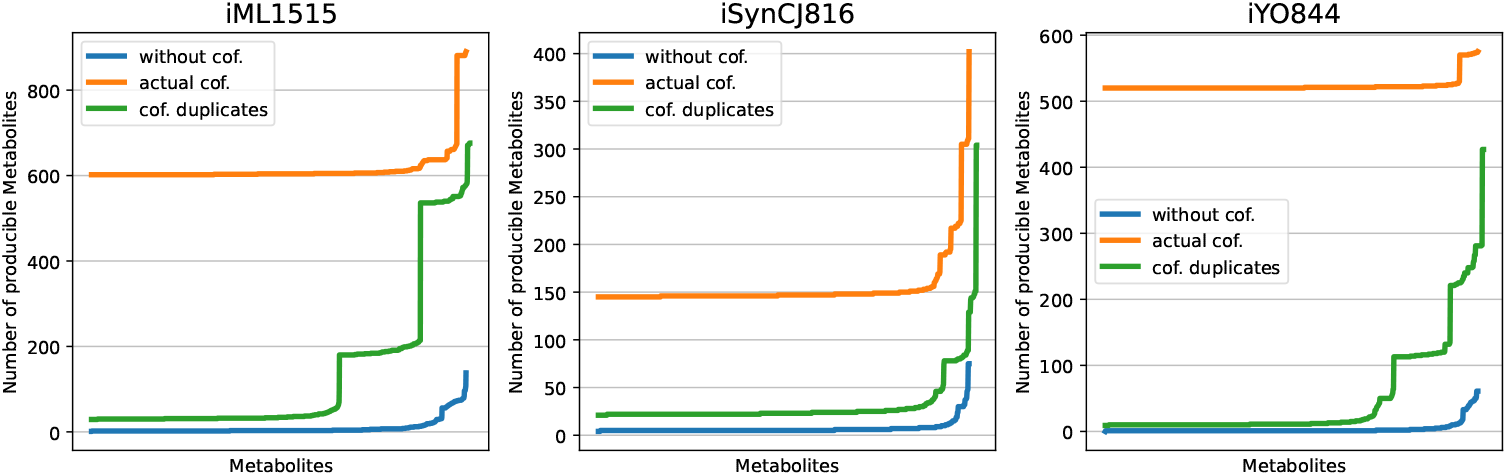
Metabolic scopes in established models of *E. coli* (iML1515), *B. subtilis* (iYO844) and *Synechocystis sp*. PCC 6803 (iSynCJ816). The differently colored graphs represent the same analysis but including no cofactors, actual cofactors and cofactor duplicates in the seed.

### Applying metabolic network expansion to a model of E. coli core metabolism

We illustrate `moped`’s topological analysis functions by applying the metabolic network expansion algorithm to a compact network of *E. coli* core metabolism, which is freely available in SBML format from the BiGG database [30]. After importing the SBML file into `moped`, we applied cofactor and reversibility duplications as described above.

For each metabolite in the network, we calculate the scope size, i.e. how many new compounds are producible if only this metabolite, water and a set of mock cofactors are available. The results of that analysis are displayed in Figure 6. In this relatively small metabolic network (72 metabolites and 95 reactions), eleven key compounds, which are mostly part of central metabolism, exhibit a largest observed scope size of 47. Such sdetailed topological analysis is useful to provide insight about central metabolites, as well as structural and functional characteristics of metabolic networks [13]. Whereas we here only display the scope size, the methods implemented in `moped` allow a far wider spectrum of analysis methods, including determination the set of producible metabolites, as well as following each step of the expansion process. The code used to produce the results and Figure 6 can be found on https://gitlab.com/marvin.vanaalst/moped-publication-2021.

**Figure 6.**
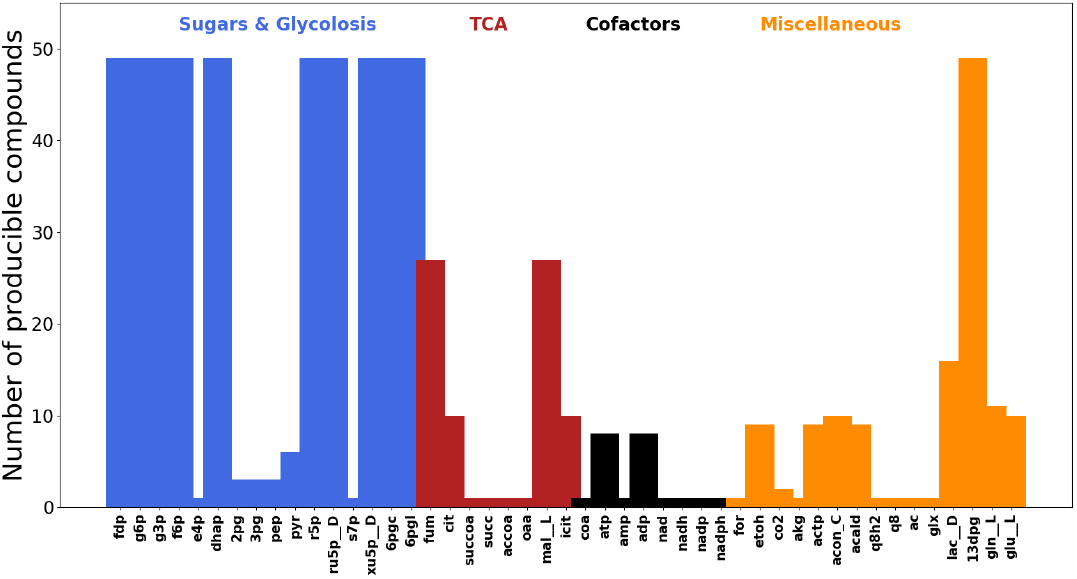
Metabolic scopes of all compounds in the *E. coli* core metabolic model calculated using moped. The Y-axis indicates the total amount of compounds producible from every compound, water and a set of acceptor mock cofactors.

### Comparison of draft reconstructions with established models and softwares

We demonstrate how `moped` provides a complete and easy-to-use pipeline to con-struct genome scale models from genome and proteome sequences and how these models can be directly applied for constraint-based analyses. For this, we download the freely available proteome sequences of *Escherichia coli* str. K-12 substr. MG1655, *Synechocystis sp*. PCC 6803 and *Bacillus subtilis* strain 168 [31–33]. We import the MetaCyc PGDB to construct a `moped` object of the MetaCyc database as a reference network. Applying the BLAST wrapper, which was described above, to the FASTA files and the reference network, we obtained three `moped` objects, representing the draft network reconstructions. Then we applied gap-filling to ensure that the reconstructed models can produce all basic biomass compounds (inspired by the *E. coli* biomass reaction from iJO1366 [34], including all nucleic acids, amino acids and lipid precursors) from M9 minimal glucose medium. For this analysis, we directly accepted all resulting gap-filling reactions. For a more accurate reconstruction, the proposed gap-filling reactions should be manually inspected before addition to the draft model. We added exchange reactions for all medium compounds, and tested if the draft models can exhibit a stationary flux distribution to produce biomass, as determined by Flux Balance Analysis. The construction of these models can be reproduced using the notebooks provided on our accompanying git.

In order to test the quality of our draft models, we compared them with established models for the respective organisms (iML1515, iYO844 and iSynCJ816) [31–33]. Furthermore, we used the same dataset and medium to construct draft models with the established genome scale modelling reconstruction software CarveMe [35]. In order to quantitatively compare all three versions of the organism network reconstructions, we used metabolic model testing (MEMOTE) pipeline to establish a fair and reproducible comparison [36]. MEMOTE calculates scores for genome scale metabolic models to evaluate the stoichiometric consistency, the GPR rules and the quality of annotations for reactions and metabolites in the respective models. A summary of the MEMOTE evaluations for the three models for the three organisms is presented in Figure 7. The MEMOTE evaluation shows that the stoichiometric consistency of draft models produced by `moped` is always of high quality. Figure 7 shows that draft models reconstructed by CarveMe and `moped` display generally good overall scores and annotations. While CarveMe draft model reconstructions show the tendency to provide better reaction annotations, `moped` draft model reconstructions display a generally better annotation of genes and GPR rules. This result shows that draft model reconstructions made with `moped` exhibit a high quality which is able to keep up with the quality of established models and software tools.

**Figure 7.**
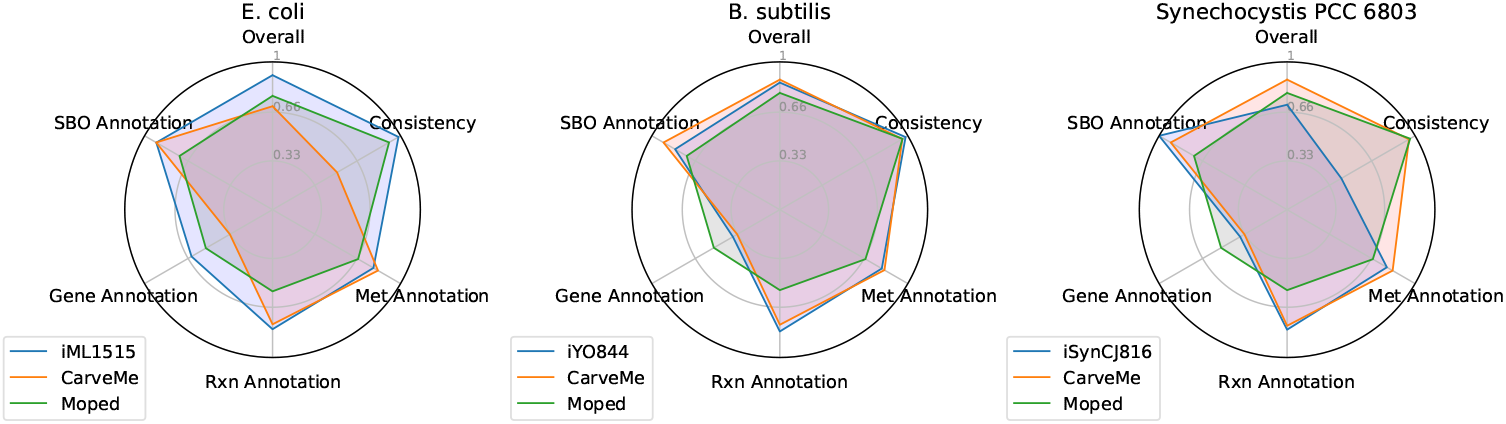
MEMOTE evaluations for draft model reconstructions produced by CarveMe and moped, as well as established models, for *E. coli, Bacillus subtilis, Synechocystis sp*. PCC 6803. MEMOTE evaluations include the stoichiometric consistency and the annotation level of models.

## Conclusion

Here, we present `moped`, a Python package representing a hub connecting the construction, modification and curation of genome scale metabolic networks with various analysis methods, which support studies of metabolic networks. `moped` supports the de novo construction of metabolic networks by importing databases, providing homology searches, including GPR associations and integrating an established gap-filling routine without the need to change software environments. Existing models from external sources can be imported using the standardized format SBML. Metabolic network models are represented as `moped` objects, which can be modified by easy-to-use and intuitive methods. `moped` models can be exported into various formats, thus integrating a diverse set of established analysis tools. Topological analysis and constraint-based optimization can be easily performed for any model represented as a `moped` object.

Examination of moped draft model reconstructions using MEMOTE demonstrated that the resulting models are generally of a high quality. The strength of draft model reconstructions with moped is the direct integration into the Python programming language: Every decision in the automatic and manual reconstruction process is documented in executable Python scripts. Therefore, the whole reconstruction process becomes fully transparent and is easily reproducible by any interested user.

The modular architecture of the open source package `moped` is particularly designed for allowing further extensions to enhance its functionality, such as the integration of additional software tools. We provide an extensive documentation for `moped`, as well as troubleshooting guides, unit-tests for all provided methods and example notebooks illustrating the usage of `moped` at https://gitlab.com/marvin.vanaalst/moped-publication-2021.

## Author Contributions

conceptualization, N.S.; implementation, N.S., M.v.A.; writing–original draft preparation, N.S.; writing–review and editing, N.S., M.v.A., and O.E.

## Data Availability Statement

Operating systems: Linux, OS X

Programming language: Python

License: GPLv3

Any restrictions to use by non-academics: For non-profit use only All source code including scripts to produce all manuscript figures can be found at https://gitlab.com/marvin.vanaalst/moped and https://gitlab.com/marvin.vanaalst/moped-publication-2021

## Acknowledgments

We thank Anna Matuszyn’ska, St. Elmo Wilken and Ovidiu Popa for critically reading our manuscript draft, Clémence Frioux for providing support for Meneco integration, Dipali Singh for giving valuable advice on MetaCyc and PDGB issues.

## Abbreviations

The following abbreviations are used in this manuscript:

PGDB: Pathway/Genome Database
FBA: Flux Balance Analysis
GPR: Gene-Protein-Reaction
SBML: Systems Biology Markup Language
ODE: Ordinary Differential Equations

## References

1. Rapoport, T.A.; Heinrich, R.; Jacobasch, G.; Rapoport, S. A linear steady-state treatment of enzymatic chains. A mathematical model of glycolysis of human erythrocytes. Eur J Biochem 1974, 42, 107–120.

2. Zomorrodi, A.R.; Segrè, D. Genome-driven evolutionary game theory helps understand the rise of metabolic interdependencies in microbial communities. Nature communications 2017, 8, 1–12.

3. Hartman, H.B.; Fell, D.A.; Rossell, S.; Jensen, P.R.; Woodward, M.J.; Thorndahl, L.; Jelsbak, L.; Olsen, J.E.; Raghunathan, A.; Daefler, S.; others. Identification of potential drug targets in Salmonella enterica sv. Typhimurium using metabolic modelling and experimental validation. Microbiology 2014, 160, 1252–1266.

4. Heinrich, R.; Schuster, S. The regulation of cellular systems; Chapman and Hall: New York, 1996.

5. Caspi, R.; Billington, R.; Keseler, I.M.; Kothari, A.; Krummenacker, M.; Midford, P.E.; Ong, W.K.; Paley, S.; Subhraveti, P.; Karp, P.D. The MetaCyc database of metabolic pathways and enzymes-a 2019 update. Nucleic acids research 2020, 48, D445–D453.

6. Karp, P.D.; Billington, R.; Caspi, R.; Fulcher, C.A.; Latendresse, M.; Kothari, A.; Keseler, I.M.; Krummenacker, M.; Midford, P.E.; Ong, Q.; others. The BioCyc collection of microbial genomes and metabolic pathways. Briefings in bioinformatics 2019, 20, 1085–1093.

7. King, Z.A.; Lu, J.; Dräger, A.; Miller, P.; Federowicz, S.; Lerman, J.A.; Ebrahim, A.; Palsson, B.O.; Lewis, N.E. BiGG Models: A platform for integrating, standardizing and sharing genome-scale models. Nucleic acids research 2016, 44, D515–D522.

8. Kanehisa, M.; Goto, S. KEGG: kyoto encyclopedia of genes and genomes. Nucleic acids research 2000, 28, 27–30.

9. Raman, K.; Chandra, N. Flux balance analysis of biological systems: applications and challenges. Briefings in bioinformatics 2009, 10, 435–449.

10. Orth, J.D.; Thiele, I.; Palsson, B.ø. What is flux balance analysis? Nature biotechnology 2010, 28, 245–248.

11. Wunderlich, Z.; Mirny, L.A. Using the topology of metabolic networks to predict viability of mutant strains. Biophysical journal 2006, 91, 2304–2311.

12. Handorf, T.; Ebenhöh, O.; Heinrich, R. Expanding metabolic networks: scopes of compounds, robustness, and evolution. Journal of molecular evolution 2005, 61, 498–512.

13. Ebenhöh, O.; Handorf, T. Functional classification of genome-scale metabolic networks. EURASIP J Bioinform Syst Biol 2009, p. 570456. doi:10.1155/2009/570456.

14. Christian, N.; May, P.; Kempa, S.; Handorf, T.; Ebenhöh, O. An integrative approach towards completing genome-scale metabolic networks. Mol Biosyst 2009, 5, 1889–1903. doi:10.1039/B915913b.

15. Orth, J.D.; Palsson, B.ø. Systematizing the generation of missing metabolic knowledge. Biotechnology and bioengineering 2010, 107, 403–412.

16. Altschul, S.F.; Gish, W.; Miller, W.; Myers, E.W.; Lipman, D.J. Basic local alignment search tool. Journal of molecular biology 1990, 215, 403–410.

17. Prigent, S.; Frioux, C.; Dittami, S.M.; Thiele, S.; Larhlimi, A.; Collet, G.; Gutknecht, F.; Got, J.; Eveillard, D.; Bourdon, J.; others. Meneco, a topology-based gap-filling tool applicable to degraded genome-wide metabolic networks. PLoS computational biology 2017, 13, e1005276.

18. Ebrahim, A.; Lerman, J.A.; Palsson, B.O.; Hyduke, D.R. COBRApy: COnstraints-based reconstruction and analysis for python. BMC systems biology 2013, 7, 74.

19. van Aalst, M.; Ebenhöh, O.; Matuszyńska, A. Constructing and analysing dynamic models with modelbase v1. 2.3: a software update. BMC bioinformatics 2021, 22, 1–15.

20. Hucka, M.; Finney, A.; Sauro, H.M.; Bolouri, H.; Doyle, J.C.; Kitano, H.; Arkin, A.P.; Bornstein, B.J.; Bray, D.; Cornish-Bowden, A.; others. The systems biology markup language (SBML): a medium for representation and exchange of biochemical network models. Bioinformatics 2003, 19, 524–531.

21. Karp, P.D.; Paley, S.; Romero, P. The pathway tools software. Bioinformatics 2002, 18, S225–S232.

22. Karpe, P.D.; Latendresse, M.; Caspi, R. The pathway tools pathway prediction algorithm. Standards in genomic sciences 2011, 5, 424–429.

23. Machado, D.; Herrgård, M.J.; Rocha, I. Stoichiometric representation of gene–protein–reaction associations leverages constraint-based analysis from reaction to gene-level phenotype prediction. PLoS computational biology 2016, 12, e1005140.

24. Thiele, I.; Palsson, B.ø. A protocol for generating a high-quality genome-scale metabolic reconstruction. Nature protocols 2010, 5, 93.

25. Kumar, V.S.; Dasika, M.S.; Maranas, C.D. Optimization based automated curation of metabolic reconstructions. BMC bioinformatics 2007, 8, 212.

26. Vitkin, E.; Shlomi, T. MIRAGE: a functional genomics-based approach for metabolic network model reconstruction and its application to cyanobacteria networks. Genome biology 2012, 13, R111.

27. Monk, J.M.; Lloyd, C.J.; Brunk, E.; Mih, N.; Sastry, A.; King, Z.; Takeuchi, R.; Nomura, W.; Zhang, Z.; Mori, H.; others. i ML1515, a knowledgebase that computes Escherichia coli traits. Nature biotechnology 2017, 35, 904–908.

28. Oh, Y.K.; Palsson, B.O.; Park, S.M.; Schilling, C.H.; Mahadevan, R. Genome-scale reconstruction of metabolic network in Bacillus subtilis based on high-throughput phenotyping and gene essentiality data. Journal of Biological Chemistry 2007, 282, 28791–28799.

29. Joshi, C.J.; Peebles, C.A.; Prasad, A. Modeling and analysis of flux distribution and bioproduct formation in Synechocystis sp. PCC 6803 using a new genome-scale metabolic reconstruction. Algal research 2017, 27, 295–310.

30. Orth, J.D.; Fleming, R.M.; Palsson, B.O. Reconstruction and use of microbial metabolic networks: the core Escherichia coli metabolic model as an educational guide. EcoSal plus 2010.

31. Blattner, F.R.; Plunkett, G.; Bloch, C.A.; Perna, N.T.; Burland, V.; Riley, M.; Collado-Vides, J.; Glasner, J.D.; Rode, C.K.; Mayhew, G.F.; others. The complete genome sequence of Escherichia coli K-12. science 1997, 277, 1453–1462.

32. Kaneko, T.; Sato, S.; Kotani, H.; Tanaka, A.; Asamizu, E.; Nakamura, Y.; Miyajima, N.; Hirosawa, M.; Sugiura, M.; Sasamoto, S.; others. Sequence analysis of the genome of the unicellular cyanobacterium Synechocystis sp. strain PCC6803. II. Sequence determination of the entire genome and assignment of potential protein-coding regions. DNA research 1996, 3, 109–136.

33. Kunst, F.; Ogasawara, N.; Moszer, I.; Albertini, A.; Alloni, G.; Azevedo, V.; Bertero, M.; Bessières, P.; Bolotin, A.; Borchert, S.; others. The complete genome sequence of the gram-positive bacterium Bacillus subtilis. Nature 1997, 390, 249–256.

34. Orth, J.D.; Conrad, T.M.; Na, J.; Lerman, J.A.; Nam, H.; Feist, A.M.; Palsson, B.ø. A comprehensive genome-scale reconstruction of Escherichia coli metabolism—2011. Molecular systems biology 2011, 7.

35. Machado, D.; Andrejev, S.; Tramontano, M.; Patil, K.R. Fast automated reconstruction of genome-scale metabolic models for microbial species and communities. Nucleic acids research 2018, 46, 7542–7553.

36. Lieven, C.; Beber, M.E.; Olivier, B.G.; Bergmann, F.T.; Ataman, M.; Babaei, P.; Bartell, J.A.; Blank, L.M.; Chauhan, S.; Correia, K.; others. MEMOTE for standardized genome-scale metabolic model testing. Nature biotechnology 2020, 38, 272–276.

